# Ibuprofen-mediated potential inhibition of biofilm development and quorum sensing in *Pseudomonas aeruginosa*

**DOI:** 10.1101/576447

**Authors:** Lu Dai, Tian-qi Wu, Zhong-hui Wang, Ruoyu Yuan, Ye Ding, Wen-chen Zhang, Shao-peng Chu, Shao-qing Ju, Juan Yu

## Abstract

*Pseudomonas aeruginosa* is one of the leading causes of opportunistic and hospital-acquired infections worldwide. The infection with *P. aeruginosa* is frequently linked with clinical treatment difficulties given drug resistance and abuse of antibiotics. Ibuprofen, a widely used non-steroidal anti-inflammatory drug, has been previously reported to exert antimicrobial activity, although the specific mechanism of its action requires additional investigation. Given the regulation effects on quorum sensing (QS), we hypothesized that inhibition of *P. aeruginosa* with ibuprofen is linked with the QS systems. First, we assessed the action of ibuprofen in *P. aeruginosa* by measuring CFU. The antimicrobial activity of ibuprofen was evaluated by crystal violent staining and acridine orange staining at various drug concentrations (0, 50, 75, and 100 μg/mL). Moreover, the effect of ibuprofen on different QS virulence factors, such as pyocyanin, elastase, protease, and rhamnolipids, was assessed revealing a concentration-dependent decrease (*P*<0.05). The effect of ibuprofen was confirmed by liquid chromatography/mass spectrometry analysis of 3-oxo-C_12_-HSL and C_4_-HSL production. In addition, qRT-PCR results identified significant suppression of Las and Rhl gene expression after 18 hours of treatment with ibuprofen (*P*<0.05), with the most significant suppression observed at the concentration of 75 μg/mL. Functional complementation with exogenous 3-oxo-C_12_-HSL and C_4_-HSL suggested that C_4_-HSL can recover the production of virulence factors and biofilm formation in *P. aeruginosa.* Molecular docking of ibuprofen with QS-associated proteins revealed high binding affinity. In summary, the results suggest that ibuprofen is a candidate drug for the treatment of clinical infections with *P. aeruginosa*.

## Introduction

*Pseudomonas aeruginosa*, the most common Gram-negative non-fermentative bacteria, is one of the leading causes of opportunistic and hospital-acquired infections worldwide(1). The infection can cause serious complications in patients with burns, cystic fibrosis, or in immunocompromised patients with respiratory infections, sepsis, osteomyelitis, endocarditis, and urinary tract infections (UTIs) (2). The infection with *P. aeruginosa* has been liked with difficulties in clinical treatment because of significant resistance of the bacterium to antibiotics (3), which is further aggravated by the worldwide abuse of antibiotics. Specifically, every year approximately 700,000 people die from antibiotic resistance worldwide, which can likely increase to 10 million by 2050 if no action is taken to reduce drug resistance or to develop new antibiotics. In 2017, the World Health Organization (WHO) published a list of drug-resistant bacteria threating the lives of people which needs require for which new antibiotics are critically needed. *P. aeruginosa* was listed second on the WHO list (4). Therefore, there is an urgent need to develop alternative strategies to treat *P. aeruginosa* infections.

Drug resistance in *P. aeruginosa* is mediated by complex mechanisms, wherein quorum sensing (QS) plays a significant role in virulence and pathogenicity. The system responsible for cell-to-cell communication in bacteria is mediated by QS, which relies on bacterial density to diffuse small signal molecules, N-acyl homoserine lactones (AHLs), that bind transcriptional activators to regulate gene expression. Studies report that QS controls expression of more than 300 genes involved in virulence factors synthesis, motility, phenotypic changes, biofilm formation, antibiotic resistance, and metabolic pathways that regulate stress response(5). *P. aeruginosa* has four QS systems: las, rhl, pqs, and iqs. Specifically, the lasI gene regulates the production of N-3-oxo-dodecanoyl-homoserine lactone (3-oxo-C_12_-HSL), while the products of lasR, including lasI and rhlR-rhlI system, require sufficient levels of 3-oxo-C_12_-HSL to activate virulence genes (6, 7). The rhl system produces N-butanoyl-L-homoserine lactone (C_4_-HSL) to modulate virulence gene expression (8). The pqs system regulates the production of 2-heptyl-3-hydroxy-4(1H)-quinolone or simple quinolone and has been reported to control biofilm formation and the production of virulence factors (5, 9). Finally, the iqs system produces 2-(2-hydroxyphenyl)-thiazole-4-carbaldehyde but its exact role requires additional research.

Studies have shown that QS may increase the expression of virulence factors, such as pyocyanin, elastase, proteases, and rhamnolipids, to adapt to the environmental pressures (10). Pyocyanin, a part of blue redox-active pigment called phenazine, has been reported to modulate bis-(3’,5’)-cyclic dimeric guanosine (c-di-GMP) messenger and extracellular polymeric substance (EPS) (11). Moreover, pyocyanin regulates secretion of the extracellular DNA (eDNA) to mediate biofilm development (11). The activity of elastase is predominantly regulated by the extracellular virulence factor LasB (elastase). It is a zinc metalloprotease acting as a vital virulence factor to control the infection with *P. aeruginosa* (12). Proteases are involved in the pathogenesis of acute lung injury caused by *P. aeruginosa* infection and are linked to the invasion and destruction of host tissue (13, 14). Rhamnolipids are biosurfactants that contribute to motility and biofilm development under stress conditions (15). Studies have shown that the level of virulence factors in QS-defective isolates is lower compared to wild-type isolates (16). Moreover, there is a relationship between the reduction in antibiofilm activity and increase in rhlR gene expression (17). Potential therapeutic strategies, attracting a lot of research attention, include targeting and inhibition of the QS system to treat *P. aeruginosa* infections.

Ibuprofen, a non-steroidal anti-inflammatory drug (NSAID), is one of the most popular non-prescription drugs given its high efficacy and safety. Early studies by Lee. et al discovered that ibuprofen inhibited pulmonary vasoconstriction and bronchiolar constriction in pigs infected with *P. aeruginosa* (18). Moreover, in an acute *P. aeruginosa* pulmonary infection in mice, ibuprofen decreased the recruitment of granulocytes to airways and suppressed lung inflammation (19). Additionally, ibuprofen effectively inhibited the production of inflammatory factor-leukotriene B_4_ (LTB_4_) in turn reducing lung inflammation in a rat model of chronic pulmonary infection (20). Importantly, in a randomized controlled trial, two-thirds of female patients with uncomplicated UTIs who took ibuprofen recovered without additional antibiotic treatment (21). Until more studies are done to carefully identify patients needing antibiotics, ibuprofen cannot be recommended as a stand-alone therapy for UTI patients. However, it is an attractive treatment alternative to antibiotics that does not cause drug resistance.

Other studies have reported that ibuprofen inhibits the growth of *P. aeruginosa* in a dose-dependent manner(22). Moreover, ibuprofen can slow the progression of lung deterioration in patients with cystic fibrosis (23). Although multiple studies have suggested that ibuprofen exerts antimicrobial activity against Gram-negative bacteria, Gram-positive bacteria, fungi, and viruses (24–27), the underlying mechanism of antimicrobial activity of ibuprofen against *P. aeruginosa* remains unclear. In this study, we hypothesized that the inhibition of *P. aeruginosa* induced by ibuprofen is related to the QS systems.

## Materials and methods

### Microbial strain, growth conditions and chemicals

*P. aeruginosa* PAO1 (ATCC15692) strain was cultured in Luria-Bertani (LB) medium (Sangon Biotech, China). Ibuprofen was purchased from Sigma-Aldrich (Cat# 14883, St. Louis, MO, USA) and dissolved in dimethyl sulfoxide (DMSO) (Biosharp). Bacteria were streaked from −80°C skim milk stocks onto blood agar plates and incubated at 37°C. Single colonies were incubated in LB medium, shaking at 120 rpm overnight at 37°C.

### Growth curve assessment

The cultures were diluted to 10^6^ colony-forming units (CFU)/mL as the initial concentration. The growth curve of PAO1 was determined in the presence or absence of different ibuprofen concentrations (0, 50, 75, and 100 μg/mL). The final DMSO concentration in all of the samples was 0.2% (v/v). All of the cultures at all of the time points (12 h, 18 h, and 24 h) were inoculated in the plates after dilution with sterile phosphate-buffered saline (PBS). The cell numbers of all of the groups were analyzed by the CFU plate count (22).

### Biofilm formation assay

PAO1 was cultured in 96-well plates in the presence or absence of different ibuprofen concentrations (0, 50, 75, and 100 μg/mL) for 24 h at 37°C. The cultures were removed and the plates were washed twice with PBS. The resulting biofilm was stained with 0.5% crystal violet for 15 min and solubilized in 95% alcohol. The absorbance was measured at 570 nm (28).

### Microscopic assessment of adherence

PAO1 was incubated in 24-well plates in the presence or absence of ibuprofen with coverslips for 24 h at 37°C. The coverslips were washed with PBS and stained with 4 μL of 0.01% acridine orange (AO, Cat#A6014, Sigma) for 10 min in the dark. The samples were immediately observed under the fluorescent microscope (Olympus BX41, Japan) at 40× lens magnification (28).

### Evaluation of virulence factor production

We used four different assays to assess the levels and activity of (1) pyocyanin, (2) elastase, (3) protease, and (4) rhamnolipids. Specific details for each assay are outlined below.

1. Pyocyanin assay: PAO1 was cultured in the presence or absence of ibuprofen at 37°C. The supernatants were collected at different times (12h, 18h, 24h), extracted with 3 mL of chloroform, and subsequently mixed with 1 mL of 0.2 N HCl. The absorbance was measured at 520 nm.
2. Elastase activity: The elastase activity was determined by Elastin-Congo red (ECR, Sigma) test according to previously published methods (29, 30). Briefly, 600 μL of supernatant was mixed with 200 μL of ECR buffer (20 mg/mL in 0.1M Tris-HCl at pH 8 with 1 mM CaCl_2_) and the mixture was incubated under shaking conditions (200 rpm) for 15 h at 37°C. Next, 100 μL of 0.12 M EDTA was used to stop the reaction. The undissolved ECR was removed by centrifugation at 5000 ×g at 10 min and the absorbance was measured at 495 nm.
3. Protease activity: The activity of protease was determined with 2% azocasein (Sigma) (30). Equal volumes of azocasein dissolved in PBS and supernatant were mixed and the cultures were incubated for 4 h at 4°C. Next, 500 μL of 10% trichloroacetic acid was added to stop the reaction and the cultures were spun at 10,000 ×g for 10 min. Finally, 500 μL of NAOH was mixed with the suspension and the absorbance was measured at 440 nm.
4. Rhamnolipid assay: Rhamnolipids were extracted from the supernatant by ethyl acetate using equal volumes (30). The samples were vortexed for 10 min and spun at 10,000 ×g for 5 min at 4°C. The upper layer was collected and the extraction process with ethyl acetate was repeated three times. The collected liquid was purified under N_2_ gas evaporation system (Agela, USA). Finally, 900 μL of orcinol regent (0.19% orcinol in 53% H_2_SO_4_) was added to the precipitate and the samples were incubated for 30 min at 80°C and the absorbance was measured at 420 nm.

### High-performance liquid chromatography with triple quadrupole mass spectrometry analysis of 3-oxo-C12-HSL and C4-HSL

The extraction method of AHL is similar to that for rhamnolipids. Based on previous reports, we used 200 μL of acetonitrile (ACN) with 0.1% formic acid to reconstitute AHLs (28, 30). Samples were injected through the C_18_ reversed-phase column (3.5 μm, 2.1 mm × 150 mm) (Waters, Milford, MA USA). The samples were analyzed using Shimadzu Nexera X2 HPLC system (Shimadzu Corporation, Japan) conjugated to AB Sciex 55000 triple quadrupole mass spectrometer (AB Sciex, Redwood City, CA USA). The flow rate of system was set to 0.3 mL/min with 0.1% (v/v) formic acid in water (as mobile phase A) and 0.1% (v/v) formic acid in ACN (as mobile phase B). The mobile phase gradient was increased to 60% B for the first 2 min after the column was equilibrated. Then the gradient was ramped to 100% B for up to 5 min and maintained for 7 min. Afterwards, the gradient was dropped to 50% B and the process was stopped at 12 min. The volume of the system was 5 μL and the column temperature was maintained at 37°C. The conditions were as follows: capillary voltage, 4000V; temperature, 350°C; GS1, 40 psi; GS2: 40 psi; collision energy, 15 eV. Analysis was conducted with the following required parameters: *m*/*z* 298.2/197 for 3-oxo-C_12_-HSL and *m*/*z* 172.0/102.1 for C_4_-HSL. The Analyst software (v. 1.2.1, AB Sciex) was used to create the calibration curve based on peak areas and analyze the data.

### Gene expression analysis by quantitative real-time PCR (qRT-PCR)

Total RNA was extracted from the cultures using the TRIzol method (TRIzol reagent, Invitrogen, Carlsbad, CA USA) and was reverse-transcribed into cDNA using a kit according to the manufacturer’s instructions (ThermoFisher Scientific, Waltham, MA, USA). The concentration and purity of RNA was determined using the UV spectrophotometer (Implen, Munchen, Germany). The process of qRT-PCR was performed by the LightCycler^®^ 480 SYBR Green I Master (Roche, Germany) in a final reaction volume of 20 μL. The expression of target genes was normalized to the expression of 16S used as a housekeeping gene. Primer sequences are listed in **Table 1**.

**Table 1.**
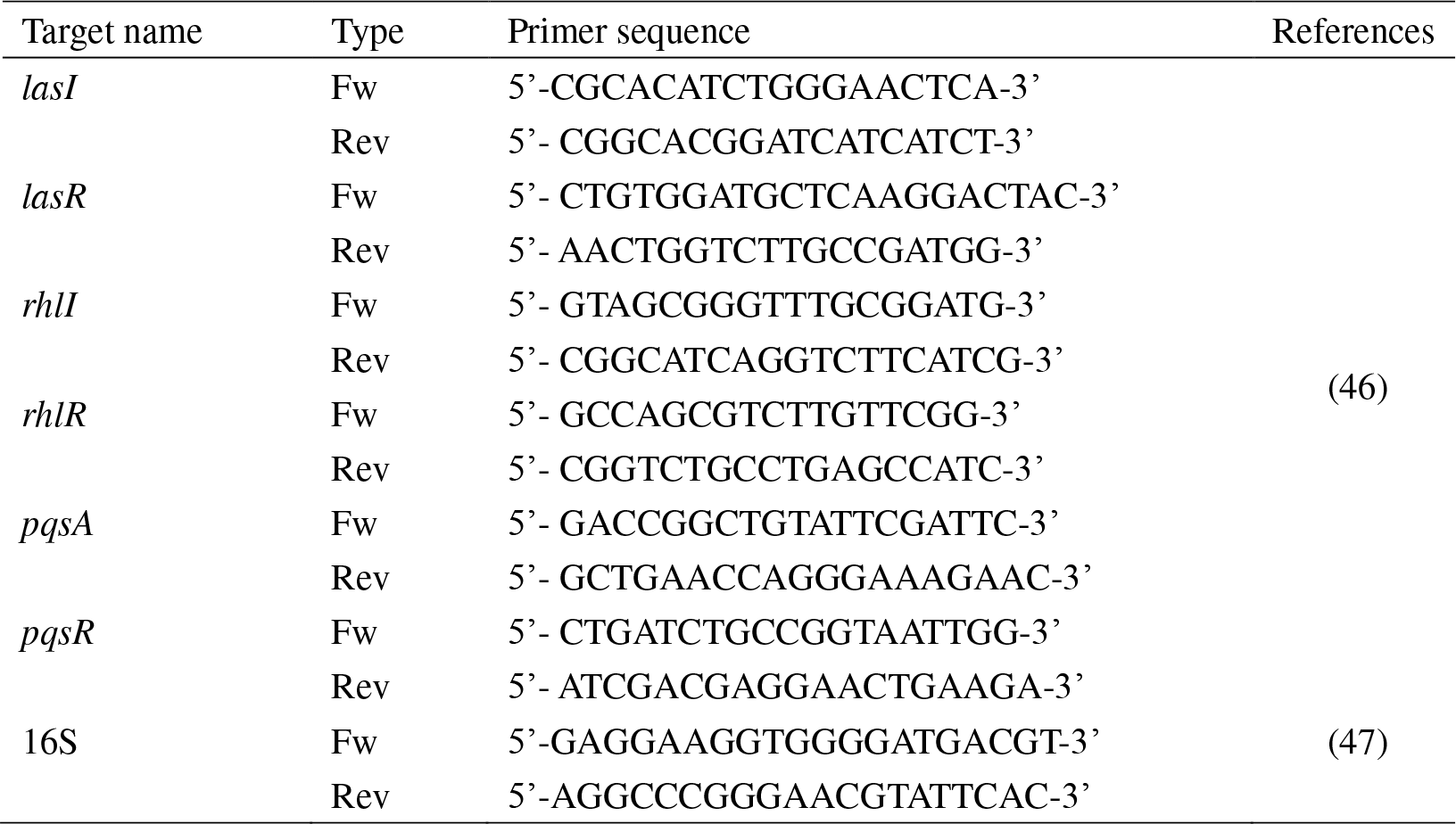
List of primers used in qRT-PCR.

### Binding analysis of ibuprofen and QS proteins

The binding interaction between the ibuprofen and proteins associated with *P. aeruginosa* QS was performed by AutoDock (31). Crystal protein structures (LasR, LasI, LuxR, LasA, pqsA, and pqsR) were downloaded from PDB (http://www.pdb.org). Since there was no available structure for RhlR, we constructed it based on the homology model using Modeller 9.17 (32). Structure of ibuprofen was drawn in ChemBioDraw (v.14) software. The structures of ibuprofen and proteins were analyzed using AutoDockTools (v. 1.5.6) and docking was carried out by AutoDock Vina.

### Functional complementation assays with 3-oxo-C12-HSL and C4-HSL

Exogenous 3-oxo-C_12_-HSL and C_4_-HSL at 2 μM concentration were added to the PAO1 medium with ibuprofen in order to carry out functional complementation assays (33). The virulence factors were determined according to methods described above.

### Statistical analysis

All of the experiments were conducted in triplicate and data are expressed as mean ± SD. The data were analyzed by one-way analysis of variance (ANOVA) using SPSS 20.0 software and the threshold for significance was set at a *p-*value< 0.05.

## Results

### Ibuprofen inhibits *P. aeruginosa* growth and biofilm formation

The bacterial burden showed a concentration-dependent reduction with increasing concentrations of ibuprofen (0, 50, 75, and 100 μg/mL) at all examined time points (12 h, 18 h, and 24h). Exposure to 50 μg/mL ibuprofen for 12 h resulted in a decrease in PAO1 count by nearly 0.7log10 compared with the control (Figure 1.A). Importantly, the reduction in bacterial count increased to approximately 2log10 when the concentration of ibuprofen was increased to 75 and 100 μg/mL (Figure 1.A). All of the cultures treated with 100 μg/mL ibuprofen had the lowest CFU counts at all of the examined time points. Since ibuprofen effectively inhibited bacterial growth, we hypothesized that ibuprofen exerts specific antibiofilm action against *P. aeruginosa*. Antibiofilm activity of ibuprofen was evaluated at several concentrations of ibuprofen by crystal violet staining. The results revealed that ibuprofen partly attenuated biofilm formation in *P. aeruginosa*, with the maximum reduction in biofilm formation (55%) found at 100 μg/mL (Figure 1.B). Next, the fluorescent microscopy was used to confirm the antibiofilm activity of ibuprofen. Specifically, *P. aeruginosa* was treated with ibuprofen and then AO was used for staining. Captured images indicated significant decrease in the attachment of biofilm cells at 100 μg/mL ibuprofen and the changes reflected a concentration-dependent pattern (Figure 1.C). These data clearly suggest that ibuprofen effectively inhibits growth and biofilm development in a concentration-dependent manner.

**Figure 1.**
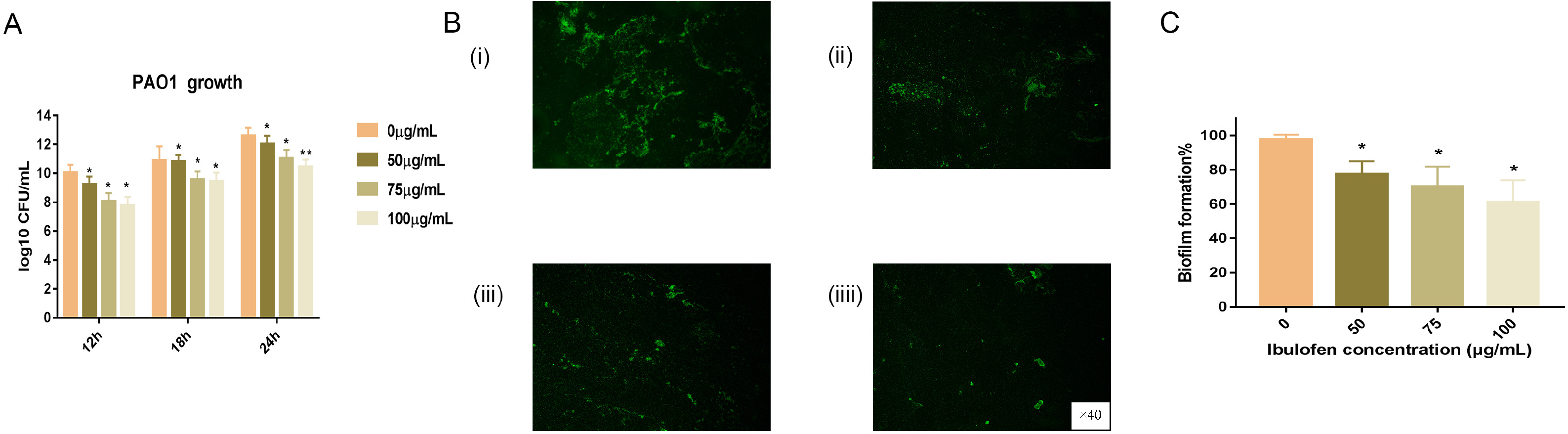
Concentration-dependent effects of ibuprofen on inhibiting the growth and biofilm formation in *P. aeruginosa*. (A) Growth conditions. (B) Adherence ability. a. 0 μg/mL ibuprofen; b. 50 μg/mL ibuprofen; c. 75 μg/mL ibuprofen; d. 100 μg/mL ibuprofen (original magnification, ×40). (C) Biofilm formation. *P. aeruginosa* PAO1 treated with different concentrations of ibuprofen (0, 50, 75, and 100 μg/mL). Data are represented as mean of four independent experiments and presented as mean ± SD. * indicates *P*<0.05, ** indicates *P*<0.01.

### Ibuprofen reduces QS signal synthesis in*P. aeruginosa*

To evaluate the action of ibuprofen on QS signaling molecules, we carefully examined the structures of 3-oxo-C_12_-HSL and C_4_-HSL (Figure 2.A and B). The HPLC/MS analysis demonstrated that levels of 3-oxo-C_12_-HSL were lower (17.08± 0.74 ng/mL) only at 100 μg/mL ibuprofen at 12 h compared to the control group and there was no evidence of concentration-dependent changes at other examined time points (Figure 2.C). In contrast, we observed a significant (*P*<0.05) time-dependent reduction in C_4_-HSL levels (Figure 2.D). PAO1 cultures treated with ibuprofen had significantly decreased levels of C_4_-HSL (up to 132.76 ng/mL) at all time points (12 h, 18 h, and 24 h). Overall, the inhibitory effects of ibuprofen on C_4_-HSL levels were concentration-dependent, with the unexceptional changes at 18 h and 24 h with the 75 μg/mL concentration (Figure 2.C).

**Figure 2.**
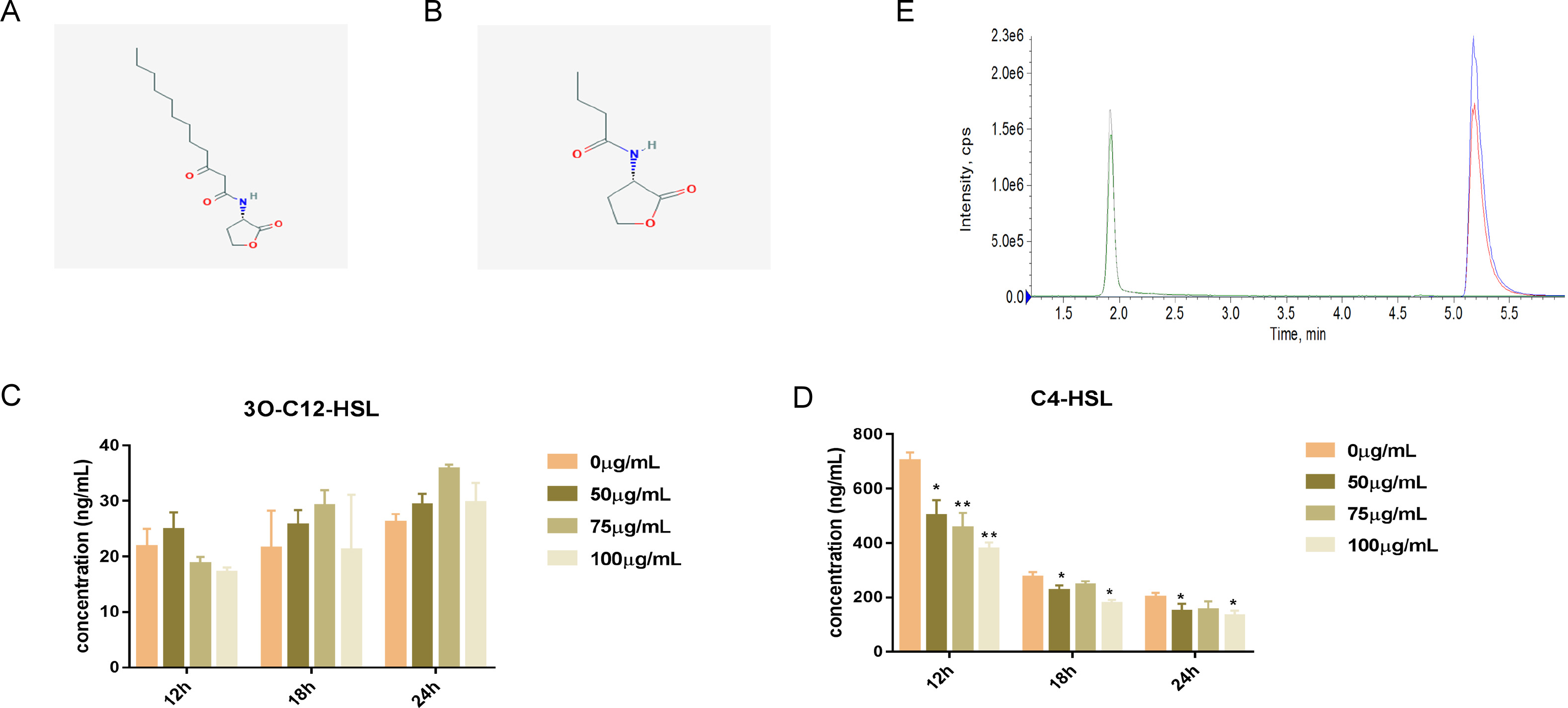
Effect of ibuprofen on AHL concentrations as determined by LC/MS. (A) Structure of 3-oxo-C_12_-HSL. (B) Structure of C_4_-HSL. (C) 3-oxo-C_12_-HSL production. (D) C_4_-HSL production. (E) LC/MS peaks of 3-oxo-C_12_-HSL and C_4_-HSL. AHL concentrations of *P. aeruginosa* PAO1 treated with different concentrations of ibuprofen (0, 50, 75, and 100 μg/mL) evaluated at three time points (12 h, 18 h, and 24 h). Data are represented as mean of three independent experiments and presented as mean ± SD. * indicates *P*<0.05, ** indicates *P*<0.01.

### Ibuprofen attenuates the production of virulence factors in *P. aeruginosa*

Since our analysis identified that ibuprofen affected bacterial signaling, we next investigated the production of virulence factors regulated in *P. aeruginosa* PAO1 strain by QS. Upon treatment, 100 μg/mL ibuprofen significantly (*P*<0.05) reduced the release of pyocyanin (up to 24.5%) at all of the examined time points (Figure 3.A). Interestingly, the significant reduction in rhamnolipid production by ibuprofen was only seen at 18 h (26.2%) and 24 h (34.5%) (Figure 3.B). There were no significant effects of ibuprofen on rhamnolipid levels at 12 h after treatment (Figure 3.B). Similarly, the production of protease was effectively inhibited by ibuprofen at 12 h and 24 h and showed a concentration-dependent effect (Figure 3.C). In contrast, only moderate, but significant (*P*<0.05), reduction (16.9%) in the total elastase production (Figure 3.D) was observed at 100 μg/mL ibuprofen at 24 h. Overall, ibuprofen exerted a significant inhibitory effect on the virulence factor production at 24 h.

**Figure 3.**
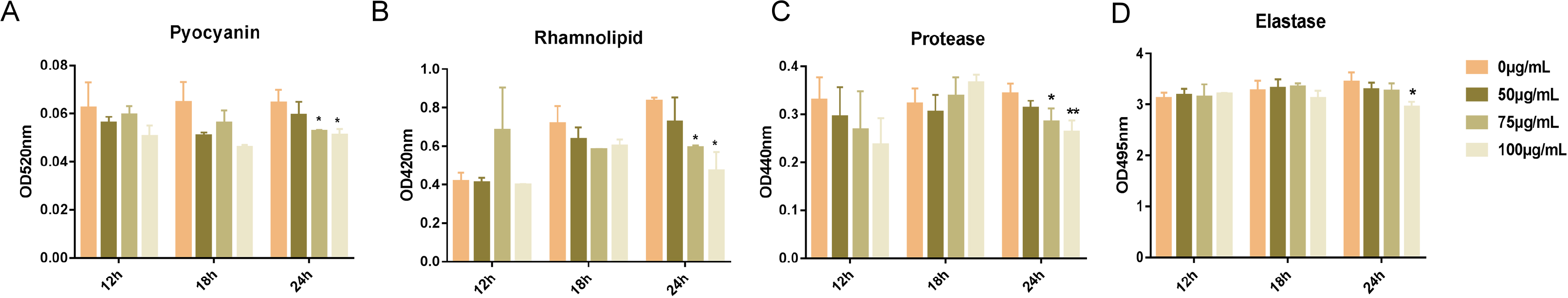
Extracellular virulence factor levels in cell-free culture supernatants treated with ibuprofen. (A) Pyocyanin. (B) Rhamnolipids. (C) Protease. (D) Elastase. *P. aeruginosa* PAO1 treated with different concentrations of ibuprofen (0, 50, 75, and 100 μg/mL) and extracellular virulence factors in the supernatants determined at three time points (12 h, 18 h, and 24 h). Data are represented as mean of three independent experiments and presented as mean ± SD. * indicates *P*<0.05, ** indicates *P*<0.01.

### The expression of QS genes decreases after ibuprofen treatment

To examine whether ibuprofen affected changes in *P. aeruginosa* on a molecular level, we analyzed gene expression levels associated with QS system by qRT-PCR (Figure 4). We noted significant (*P*<0.05) reduction in the expression of lasI, lasR, rhll, rhlR, pqsA, and pqsR at 18 h. However, the effect did not follow a concentration-dependent pattern, like in the case of virulence factor assessment. Among all of the examined concentrations, 75 μg/mL exerted the most pronounced effects in suppressing QS system gene expression. The relative expression of lasI and lasR was decreased 1.5- and 3.5-fold, respectively (Figure 4.A and B), while the expression of pqsA and pqsR was reduced by approximately 1.5~2.0-fold (Figure 4.C and D). Most interesting, the levels of rhlI were reduced 4.7-fold and the levels of rhlR was suppressed 8.3-fold (Figure 4.E and F). These data suggest that ibuprofen mainly acted on the Rhl and Las system.

**Figure 4.**
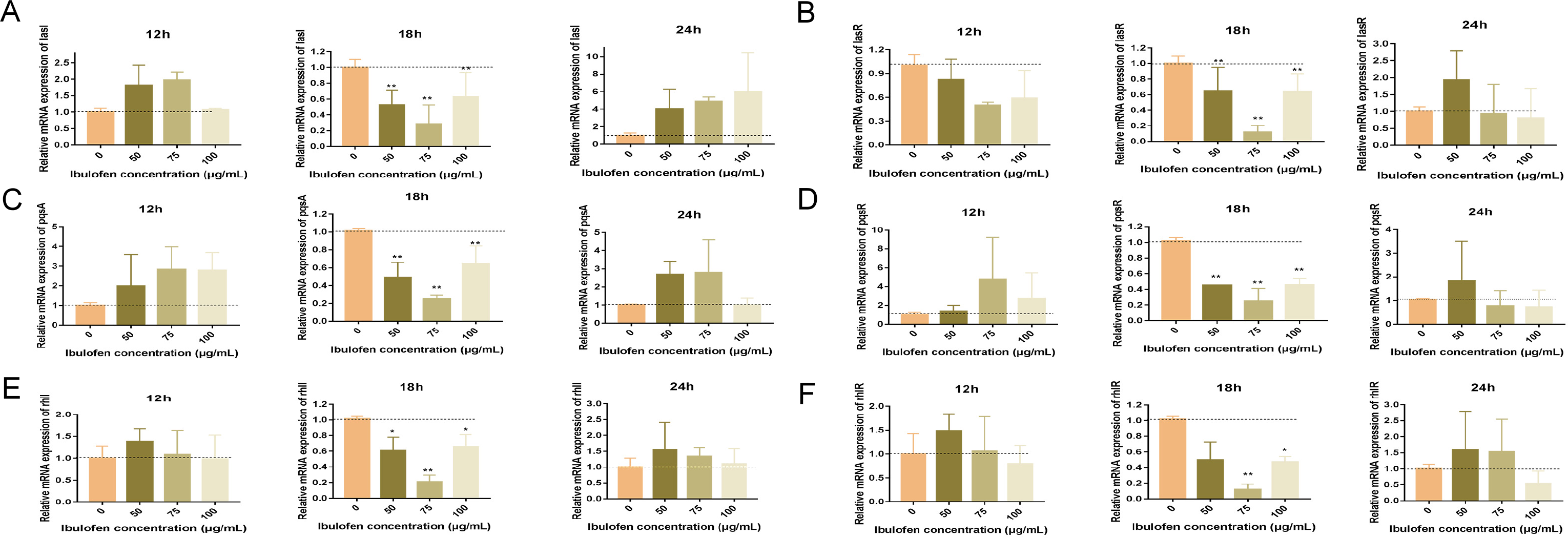
The changes in gene expression related to the QS system in *P. aeruginosa* treated with ibuprofen. (A) lasI. (B) lasR. (C) pqsA. (D) pqsR. (E) rhlI. (F) rhlR. *P. aeruginosa* PAO1 was treated with different concentrations of ibuprofen (0, 50, 75, 100 μg/mL) and the gene expression changes were evaluated at three time points (12 h, 18 h, and 24 h). Data are represented as mean of three independent experiments and presented as mean ± SD. * indicates *P*<0.05, ** indicates *P*<0.01.

Given the inhibition of QS signaling molecules, we examined whether exogenous addition of 3-oxo-C_12_-HSL and C_4_-HSL could functionally complement the activity of ibuprofen. Specifically, we observed a significant increase (*P*<0.05) in biofilm formation (Figure 5.A), pyocyanin production (Figure 5.B), elastase activity (Figure 5.C), protease activity (Figure 5.D), and rhamnolipid production (Figure 5.E) upon addition of exogenous C_4_-HSL. Importantly, we did not observe any statistically significant changes in the level of virulence factors upon treatment with exogenous 3-oxo-C_12_-HSL (Figure 5).

**Figure 5.**
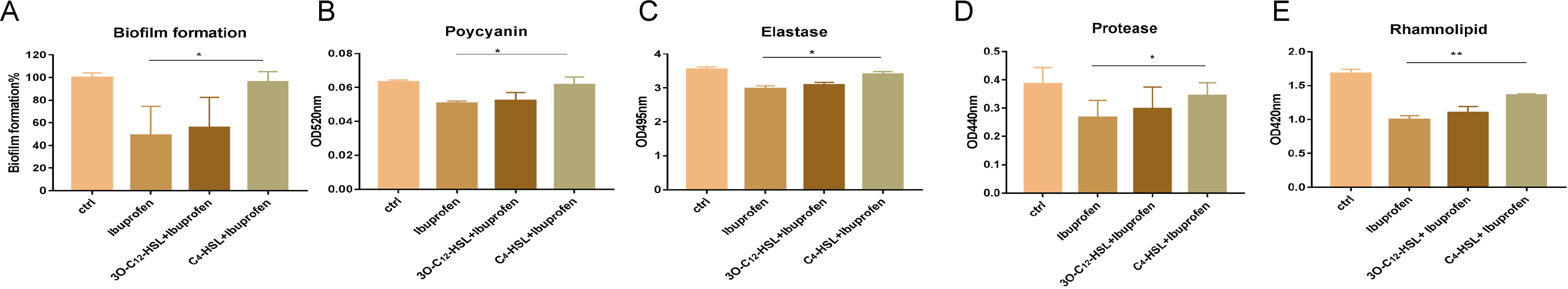
Functional complementation of ibuprofen-treated *P. aeruginosa* with exogenous 3-oxo-C_12_-HSL and C_4_-HSL. (A) Biofilm formation. (B) Pyocyanin. (C) Elastase. (D) Protease. (E) Rhamnolipids. The determination of virulence factor production and biofilm formation in *P. aeruginosa* PAO1 treated with ibuprofen (100 μg/mL) in the presence of 3-oxo-C_12_-HSL and C_4_-HSL. Data are represented as mean of three independent experiments and presented as mean ± SD. * indicates *P*<0.05, ** indicates *P*<0.01.

### Assessment of molecular docking of ibuprofen with QS-related proteins in *P. aeruginosa*

To further analyze the role of ibuprofen, molecular docking study was performed to better understand the effect of ibuprofen on attenuating QS-related proteins (LasA, LasI, RhlR, LuxR, LasR, PqsA, and PqsR). Crystal structures of the QS-related proteins were found online: LasR (PDB ID: 2UV0), LasI (PDB ID: IRO5), LuxR (PDB ID: 3JPU), LasA (PDB ID: 3IT7), PqsA (PDB ID:5OE3), and PqsR (PDB ID:5OE3). Although the crystal structure of RhlR was not available, we obtained it by homology modeling. The native receptors and binding affinity of ibuprofen with QS-related proteins is summarized in **Table 2**. Specifically, we observed that ibuprofen was docked with LasA at two different binding pockets occupied by two different ligands: tartaric acid (TLA) and glycerol (GOL). The binding energy computed using AutoDock between ibuprofen and LasA was −6.7 kcal/mol with the GOL ligand (**Table 2**). Moreover, ibuprofen formed three hydrogen bonds with Ser50 of LasA at a distance of 3.0, 3.2 and 3.4 Å (Figure 6.A). In addition, ibuprofen formed a hydrogen bond with Arg12 of LasA at a distance of 3.4 Å (Figure 6.A). The results of AutoDock analysis suggested that ibuprofen only formed a hydrogen bond with Ile107 of LasI (Figure 6.B). The binding energy between ibuprofen and LasI was −7.4 kcal/mol (**Table 2**). We constructed a virtual homology model of RhlR since the crystal structure was not available. Here, ibuprofen formed two hydrogen bonds with Asp80 of RhlR at a distance of 1.5 and 3.3 Å (Figure 6.C). The docking score between ibuprofen and RhlR was −7.2 kcal/mol (**Table 2**). Moreover, ibuprofen formed two hydrogen bonds with Thr75 and Asp73 in LuxR at a distance of 3.2 and 1.9 Å, respectively (Figure 6.D). The result showed that binding energy between ibuprofen and LuxR was −7.9 kcal/mol (**Table 2**). The docking studies showed that ibuprofen binds into the active site of LasR with Asp73, Thr75, Thr115, and Ser129 residues by hydrogen bond interactions (Figure 6.E). The binding energy between ibuprofen and LasR was −7.7 kcal/mol (**Table 2**). PqsA was shown to interact with four different ligands, and 5’-O-[(S)-[(2-aminobenzoyl)oxy](hydroxy)phosphoryl]adenosine (3UK) with high binding scores (−6.8 kcal/mol) (**Table 2**). Ibuprofen formed hydrogen bonds with Asp299, Ser280, Gly279, and Gly300 residues at a distance of 1.9, 3.2, 2.5, and 3.4 Å, respectively (Figure 6.F). Furthermore, ibuprofen was found to interact with the Ser196 and Gln194 of PqsR (Figure 6.G) and showed binding energy of −6.8 kcal/mol (**Table 2**). After docking of ibuprofen into the original ligand binding pockets of these proteins, we found that ibuprofen was binding with LuxR (−7.9 kcal/mol), LasR (−7.7 kcal/mol), LasI (−7.4 kcal/mol), and RhlR (−7.2 kcal/mol) with high binding scores (**Table 2**). Thus, the molecular docking data further demonstrate that ibuprofen may inhibit the LuxR, Las, and Rhl systems in *P. aeruginosa*.

**Table 2.** Binding affinity of QS regulator proteins from *P. aeruginosa* with ibuprofen.

| Receptor protein | PDB ID | Native ligand | Probable inhibitor | Hydrogen bonds | Binding energy(kcal/mol) |
| --- | --- | --- | --- | --- | --- |
| LasA | 3IT7 | GOL | Ibuprofen | Ser50, Arg12 | -6.7 |
| LasI | 1RO5 | perrhenate | Ibuprofen | Ile107 | -7.4 |
| RhlR | 4Y15 | - | Ibuprofen | Asp80 | -7.2 |
| LuxR | 3JPU | TY4 | Ibuprofen | Thr75, Asp73 | -7.9 |
| LasR | 2UV0 | OHN | Ibuprofen | Asp73, Thr75, Thr115, Ser129 | -7.7 |
| PqsA | 5OE3 | 3UK | Ibuprofen | Asp299, Ser280, Gly279, Gly300 | -6.8 |
| PqsR | 4JVD | NNQ | Ibuprofen | Ser196, Gln194 | -6.7 |

AutoDock provides a binding score revealing the binding affinity measured in kcal/mol. Negative scores indicate high binding affinity.

**Figure 6.**
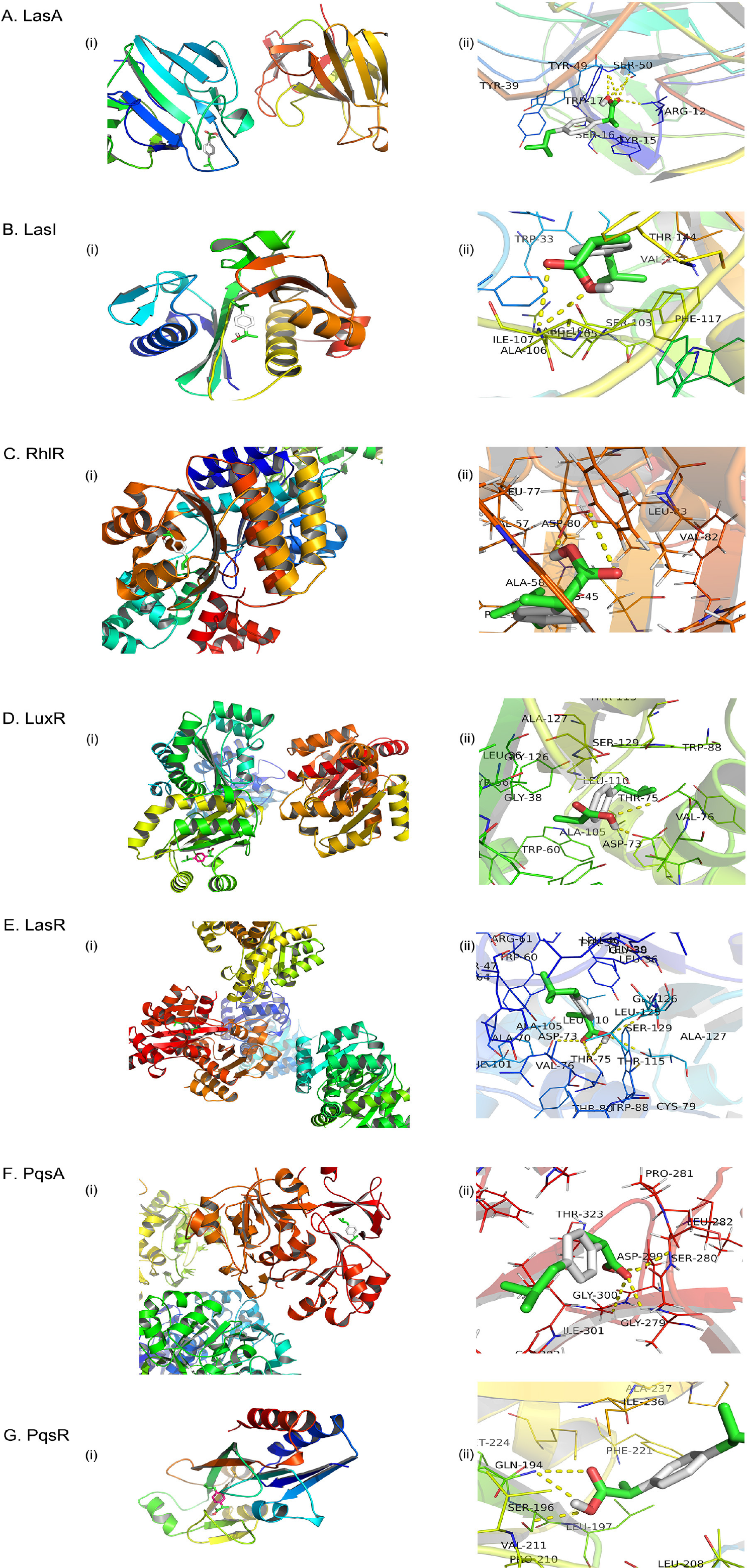
Ibuprofen docked in different binding pockets of QS proteins in *P. aeruginosa*. (A) Cartoon structure of LasA with ibuprofen. (B) Cartoon structure of LasI with ibuprofen. (C) Cartoon structure of RhlR with ibuprofen. (D) Cartoon structure of LuxR with ibuprofen. (E) Cartoon structure of LasR with ibuprofen. (F) Cartoon structure of PqsA with ibuprofen. (G) Cartoon structure of PqsR with ibuprofen. Proteins are represented in cartoons and ligands are represented in stick models.

## Discussion

Inhibitors of QS which act to directly prevent biofilm formation and limit virulence factor release are being considered as a novel strategy for treating *P. aeruginosa* infections (34–36). Although many studies examined the action of ibuprofen in bacterial infections, the antibacterial mechanism of action of ibuprofen requires further exploration. The first mention of the antifungal and antimicrobial activity of ibuprofen was reported by Hersh and colleagues in 1991. Many reports have demonstrated that ibuprofen exerts antibacterial effects in high concentrations that exceed levels seen in normal human blood (24, 37, 38). Previous report indicated that ibuprofen was able to prevent AHL inhibition of the responses to nucleotides in cystic fibrosis airway epithelium (39). Moreover, ibuprofen has been reported as an alternative drug for the treatment of local infections (e.g., UTIs) and showed a promising therapeutic potential when combined with clinical antibiotics (21, 24, 40–42).

In this study, we confirmed that ibuprofen has antimicrobial activity and specifically reduces the QS signaling molecules. Based on the clinical trial data, the efficacy and safety of high-dose ibuprofen (50-100 μg/mL plasma concentration) was shown in cystic fibrosis patients (22). Based on these data, we designed a concentration gradient of ibuprofen that was tested. We discovered that ibuprofen can affect the growth of *P. aeruginosa* but it may not have a complete bactericidal effect (Figure1.A). The OA staining assay demonstrated that the attachment activity of *P. aeruginosa* was inhibited by ibuprofen in a concentration-dependent manner (Figure 1.C). Biofilm is defined as the community of bacteria attached to both biological and abiotic surfaces, and the presence of biofilm further exacerbates the crisis of antibiotic resistance worldwide (43). Crystal violet staining confirmed that ibuprofen exerted a significant effect on the reduction of biofilm formation by *P. aeruginosa* (Figure 1.B), with the 100 μg/mL concentration having the greatest effect on biofilm inhibition. We next determined that the synthesis of C_4_-HSL was inhibited by ibuprofen in a concentration- and time-dependent manner (Figure 2.C). However, ibuprofen did not reduce 3-oxo-C_12_-HSL secretion, suggesting that ibuprofen exerts antimicrobial effects through the reduction of C_4_-HSL levels instead of the direct cell death effects (Figure 1 and 2).

Many studies have been conducted to identify that QS molecules act as regulators mediating the production of virulence factors (36, 44). Studies in different animal models demonstrate that QS plays a critical role in the pathogenicity of *P. aeruginosa* infections (10, 17, 45, 46). In this study, we found that the virulence factors, including pyocyanin, elastase, protease, and rhamnolipids, were significantly reduced upon treatment with ibuprofen at the 24 h time point in a concentration-dependent manner (Figure 3). We concluded that treatment with ibuprofen specifically and effectively targets QS molecules and virulence factors in *P. aeruginosa* and can therefore be a promising clinical therapy.

Las and Rhl are involved in the LuxR-type QS system of *P. aeruginosa*. Gene expression analysis revealed significant decreases in genes encoding QS proteins (lasI, lasR, rhlI, rhlR, pasA, and pqsR) following ibuprofen treatment within 18 h (Figure 4). Despite these findings, the inhibition by ibuprofen did not show concentration-dependent changes and future studies are needed to address this finding in detail. Work by Somaia. et al suggested that aspirin, another NSAID, reduces lasR gene expression by competing with 3-oxo-C_12_-HSL (47). In our findings, ibuprofen significantly repressed lasR and rhlR gene expression by 88.3% and 88%, respectively. The antimicrobial activity of ibuprofen against *P. aeruginosa* is dependent on lasR and rhlR. The reduction in pqsA and pqsR gene expression levels by ibuprofen was almost 75%. In our study, the decreased level of virulence factors was consistent with the reduction in QS systems related to the effects on Las and Rhl gene expression (Figure 3 and 4).

To further investigate the antimicrobial activity of ibuprofen, we examined the molecular interaction of QS-associated LasR, LasI, LuxR, LasA, RhlR, PqsA, and PqsR proteins with ibuprofen. Our docking studies of ibuprofen with proteins suggested that ibuprofen specifically binds to ligand binding pockets in four of the QS proteins LuxR, LasR, LasI, and RhlR with a relatively high binding affinity (Figure 6 and **Table 2**). Amino acids presenting in the binding sites of these four proteins play crucial roles in the dimerization and activation of the receptors. Based on our results, we propose that the binding of ibuprofen to the amino acids might be directly responsible for the inactivation of the QS proteins. We believe that this occurs either by preventing protein dimerization or there are direct effects on the DNA resulting in suppression of gene expression downstream in the QS system. Overall, the data suggest that the proteins related to QS may be a promising target for antibacterial and antibiofilm action of ibuprofen.

Moreover, we also examined whether exogenous addition of 3-oxo-C_12_-HSL and C_4_-HSL can functionally complement the activity of ibuprofen. We observed that C_4_-HSL was capable of reversing the antimicrobial action of ibuprofen on biofilm formation and virulence factors, whereas 3-oxo-C_12_-HSL had no effect (Figure 5). These data recapitulated the LC/MS results. Although 3-oxo-C_12_-HSL-LasR can directly activate transcription of target genes and stimulate the production of C_4_-HSL and RhlR, the action of ibuprofen was not inhibited by 3-oxo-C_12_-HSL. These results warrant additional investigation to further explore the specific antimicrobial role of ibuprofen.

In summary, we demonstrated the molecular mechanism of antibacterial activity of ibuprofen against *P. aeruginosa*. In our study, ibuprofen prevented the *P. aeruginosa* by directly reducing bacterial burden and suppressing the LuxR QS system. Considering the gene expression results and effect on virulence factors, treatment with ibuprofen may be a promising therapy when dealing with *P. aeruginosa* infections. Additionally, because other studies suggest that ibuprofen is effective in treating UTIs and cystic fibrosis, we propose ibuprofen as a candidate drug to be used in the management of clinical infections with *P. aeruginosa*.

## Acknowledgements

This work was supported by the Science and Technology Project of Nantong city (2018).

## References

1. Pang Z, Raudonis R, Glick BR, Lin TJ, Cheng Z. 2019. Antibiotic resistance in Pseudomonas aeruginosa: mechanisms and alternative therapeutic strategies. Biotechnol Adv 37:177–192.

2. Moradali MF, Ghods S, Rehm BH. 2017. Pseudomonas aeruginosa Lifestyle: A Paradigm for Adaptation, Survival, and Persistence. Front Cell Infect Microbiol 7:39.

3. Chevalier S, Bouffartigues E, Bodilis J, Maillot O, Lesouhaitier O, Feuilloley MGJ, Orange N, Dufour A, Cornelis P. 2017. Structure, function and regulation of Pseudomonas aeruginosa porins. FEMS Microbiol Rev 41:698–722.

4. Willyard C. 2017. The drug-resistant bacteria that pose the greatest health threats. Nature 543:15.

5. Azam MW, Khan AU. 2019. Updates on the pathogenicity status of Pseudomonas aeruginosa. Drug Discov Today 24:350–359.

6. Davies DG, Parsek MR, Pearson JP, Iglewski BH, Costerton JW, Greenberg EP. 1998. The involvement of cell-to-cell signals in the development of a bacterial biofilm. Science 280:295–8.

7. Jakobsen TH, Bjarnsholt T, Jensen PO, Givskov M, Hoiby N. 2013. Targeting quorum sensing in Pseudomonas aeruginosa biofilms: current and emerging inhibitors. Future Microbiol 8:901–21.

8. Jimenez PN, Koch G, Thompson JA, Xavier KB, Cool RH, Quax WJ. 2012. The multiple signaling systems regulating virulence in Pseudomonas aeruginosa. Microbiol Mol Biol Rev 76:46–65.

9. Lee J, Zhang L. 2015. The hierarchy quorum sensing network in Pseudomonas aeruginosa. Protein Cell 6:26–41.

10. Karatuna O, Yagci A. 2010. Analysis of quorum sensing-dependent virulence factor production and its relationship with antimicrobial susceptibility in Pseudomonas aeruginosa respiratory isolates. Clin Microbiol Infect 16:1770–5.

11. Meirelles LA, Newman DK. 2018. Both toxic and beneficial effects of pyocyanin contribute to the lifecycle of Pseudomonas aeruginosa. Mol Microbiol 110:995–1010.

12. Szamosvari D, Reichle VF, Jureschi M, Bottcher T. 2016. Synthetic quinolone signal analogues inhibiting the virulence factor elastase of Pseudomonas aeruginosa. Chem Commun (Camb) 52:13440–13443.

13. Kipnis E, Sawa T, Wiener-Kronish J. 2006. Targeting mechanisms of Pseudomonas aeruginosa pathogenesis. Med Mal Infect 36:78–91.

14. Lindsay S, Oates A, Bourdillon K. 2017. The detrimental impact of extracellular bacterial proteases on wound healing. Int Wound J 14:1237–1247.

15. Reis RS, Pereira AG, Neves BC, Freire DM. 2011. Gene regulation of rhamnolipid production in Pseudomonas aeruginosa--a review. Bioresour Technol 102:6377–84.

16. Garcia-Contreras R. 2016. Is Quorum Sensing Interference a Viable Alternative to Treat Pseudomonas aeruginosa Infections? Front Microbiol 7:1454.

17. Gholamrezazadeh M, Shakibaie MR, Monirzadeh F, Masoumi S, Hashemizadeh Z. 2018. Effect of nano-silver, nano-copper, deconex and benzalkonium chloride on biofilm formation and expression of transcription regulatory quorum sensing gene (rh1R) in drug-resistance Pseudomonas aeruginosa burn isolates. Burns 44:700–708.

18. Lee CC, Sugerman HJ, Tatum JL, Wright TP, Hirsh PD, Hirsch JI. 1986. Effects of ibuprofen on a pig Pseudomonas ARDS model. J Surg Res 40:438–44.

19. Sordelli DO, Cerquetti MC, el-Tawil G, Ramwell PW, Hooke AM, Bellanti JA. 1985. Ibuprofen modifies the inflammatory response of the murine lung to Pseudomonas aeruginosa. Eur J Respir Dis 67:118–27.

20. Konstan MW, Vargo KM, Davis PB. 1990. Ibuprofen attenuates the inflammatory response to Pseudomonas aeruginosa in a rat model of chronic pulmonary infection. Implications for antiinflammatory therapy in cystic fibrosis. Am Rev Respir Dis 141:186–92.

21. Gagyor I, Bleidorn J, Kochen MM, Schmiemann G, Wegscheider K, Hummers-Pradier E. 2015. Ibuprofen versus fosfomycin for uncomplicated urinary tract infection in women: randomised controlled trial. Bmj 351:h6544.

22. Shah PN, Marshall-Batty KR, Smolen JA, Tagaev JA, Chen Q, Rodesney CA, Le HH, Gordon VD, Greenberg DE, Cannon CL. 2018. Antimicrobial Activity of Ibuprofen against Cystic Fibrosis-Associated Gram-Negative Pathogens. Antimicrob Agents Chemother 62:1–22.

23. Lands LC, Stanojevic S. 2016. Oral non-steroidal anti-inflammatory drug therapy for lung disease in cystic fibrosis. Cochrane Database Syst Rev 4:Cd001505.

24. Obad J, Suskovic J, Kos B. 2015. Antimicrobial activity of ibuprofen: new perspectives on an “Old” non-antibiotic drug. Eur J Pharm Sci 71:93–8.

25. Ogundeji AO, Pohl CH, Sebolai OM. 2016. Repurposing of Aspirin and Ibuprofen as Candidate Anti-Cryptococcus Drugs. Antimicrob Agents Chemother 60:4799–808.

26. Pan T, Peng Z, Tan L, Zou F, Zhou N, Liu B, Liang L, Chen C, Liu J, Wu L, Liu G, Peng Z, Liu W, Ma X, Zhang J, Zhu X, Liu T, Li M, Huang X, Tao L, Zhang Y, Zhang H. 2018. Non-Steroidal Anti-Inflammatory Drugs (NSAIDs) Potently Inhibit the Replication of Zika Viruses by Inducing the Degradation of AXL. J Virol doi:10.1128/jvi.01018-18.

27. Laudy AE, Mrowka A, Krajewska J, Tyski S. 2016. The Influence of Efflux Pump Inhibitors on the Activity of Non-Antibiotic NSAIDS against Gram-Negative Rods. PLoS One 11:e0147131.

28. Chatterjee M, D’Morris S, Paul V, Warrier S, Vasudevan AK, Vanuopadath M, Nair SS, Paul-Prasanth B, Mohan CG, Biswas R. 2017. Mechanistic understanding of Phenyllactic acid mediated inhibition of quorum sensing and biofilm development in Pseudomonas aeruginosa. Appl Microbiol Biotechnol 101:8223–8236.

29. Das MC, Sandhu P, Gupta P, Rudrapaul P, De UC, Tribedi P, Akhter Y, Bhattacharjee S. 2016. Attenuation of Pseudomonas aeruginosa biofilm formation by Vitexin: A combinatorial study with azithromycin and gentamicin. Sci Rep 6:23347.

30. Jack AA, Khan S, Powell LC, Pritchard MF, Beck K, Sadh H, Sutton L, Cavaliere A, Florance H, Rye PD, Thomas DW, Hill KE. 2018. Alginate Oligosaccharide-Induced Modification of the lasI-lasR and rhlI-rhlR Quorum-Sensing Systems in Pseudomonas aeruginosa. Antimicrob Agents Chemother 62:1–14.

31. Trott O, Olson AJ. 2010. AutoDock Vina: improving the speed and accuracy of docking with a new scoring function, efficient optimization, and multithreading. J Comput Chem 31:455–61.

32. Fiser A, Sali A. 2003. Modeller: generation and refinement of homology-based protein structure models. Methods Enzymol 374:461–91.

33. Babic F, Venturi V, Maravic-Vlahovicek G. 2010. Tobramycin at subinhibitory concentration inhibits the RhlI/R quorum sensing system in a Pseudomonas aeruginosa environmental isolate. BMC Infect Dis 10:148.

34. Janssens JC, De Keersmaecker SC, De Vos DE, Vanderleyden J. 2008. Small molecules for interference with cell-cell-communication systems in Gram-negative bacteria. Curr Med Chem 15:2144–56.

35. Hardman AM, Stewart GS, Williams P. 1998. Quorum sensing and the cell-cell communication dependent regulation of gene expression in pathogenic and non-pathogenic bacteria. Antonie Van Leeuwenhoek 74:199–210.

36. Le Berre R, Faure K, Nguyen S, Pierre M, Ader F, Guery B. 2006. {Quorum sensing: a new clinical target for Pseudomonas aeruginosa?]. Med Mal Infect 36:349–57.

37. Elvers KT, Wright SJ. 1995. Antibacterial activity of the anti-inflammatory compound ibuprofen. Lett Appl Microbiol 20:82–4.

38. Pina-Vaz C, Sansonetty F, Rodrigues AG, Martinez-De-Oliveira J, Fonseca AF, Mardh PA. 2000. Antifungal activity of ibuprofen alone and in combination with fluconazole against Candida species. J Med Microbiol 49:831–40.

39. Saleh A, Figarella C, Kammouni W, Marchand-Pinatel S, Lazdunski A, Tubul A, Brun P, Merten MD. 1999. Pseudomonas aeruginosa quorum-sensing signal molecule N-(3-oxododecanoyl)-L-homoserine lactone inhibits expression of P2Y receptors in cystic fibrosis tracheal gland cells. Infect Immun 67:5076–82.

40. Little P. 2017. Antibiotics or NSAIDs for uncomplicated urinary tract infection? Bmj 359:j5037.

41. Vik I, Bollestad M, Grude N, Baerheim A, Damsgaard E, Neumark T, Bjerrum L, Cordoba G, Olsen IC. 2018. Ibuprofen versus pivmecillinam for uncomplicated urinary tract infection in women-A double-blind, randomized non-inferiority trial. PLoS Med 15:e1002569.

42. Pina-Vaz C, Rodrigues AG, Costa-de-Oliveira S, Ricardo E, Mardh PA. 2005. Potent synergic effect between ibuprofen and azoles on Candida resulting from blockade of efflux pumps as determined by FUN-1 staining and flow cytometry. J Antimicrob Chemother 56:678–85.

43. Mathur H, Field D. 2018. Fighting biofilms with lantibiotics and other groups of bacteriocins. NPJ Biofilms Microbiomes 4:9.

44. Alayande AB, Aung MM, Kim IS. 2018. Correlation Between Quorum Sensing Signal Molecules and Pseudomonas aeruginosa’s Biofilm Development and Virulency. Curr Microbiol 75:787–793.

45. Cosson P, Zulianello L, Join-Lambert O, Faurisson F, Gebbie L, Benghezal M, Van Delden C, Curty LK, Kohler T. 2002. Pseudomonas aeruginosa virulence analyzed in a Dictyostelium discoideum host system. J Bacteriol 184:3027–33.

46. Rahme LG, Stevens EJ, Wolfort SF, Shao J, Tompkins RG, Ausubel FM. 1995. Common virulence factors for bacterial pathogenicity in plants and animals. Science 268:1899–902.

47. El-Mowafy SA, Abd El Galil KH, El-Messery SM, Shaaban MI. 2014. Aspirin is an efficient inhibitor of quorum sensing, virulence and toxins in Pseudomonas aeruginosa. Microb Pathog 74:25–32.

